# Development of serum substitute medium for bone tissue engineering

**DOI:** 10.1101/2022.10.07.511271

**Authors:** Sana Ansari, Keita Ito, Sandra Hofmann

## Abstract

In tissue engineering, cells are grown often on scaffolds and subjected to chemical/mechanical stimuli. Most such cultures still use fetal bovine serum (FBS) despite its known disadvantages including ethical concerns, safety issues, and variability in composition, which greatly influences the experimental outcomes. To overcome the disadvantages of using FBS, chemically defined serum substitute medium needs to be developed. Development of such medium depends on cell type and application - which makes it impossible to define one universal serum substitute medium for all cells in any application. Here, we developed a serum substitute medium for bone tissue engineering (BTE) in a step-by-step process. Essential components were added to the medium while human bone marrow mesenchymal stromal cells (hBMSCs, osteoblast progenitor cells) were cultured in two-dimensional (2D) and three-dimensional (3D) substrates. In a 3-week culture, the developed serum substitute medium worked equally well as FBS containing medium in term of cell attachment to the substrate, cell survival, osteoblast differentiation, and deposition of extracellular matrix. In the next step, the use of serum substitute medium was evaluated when culturing cells under mechanical loading in the form of shear stress. The outcomes showed that the application of shear stress is essential to improve extracellular matrix formation while using serum substitute medium. The developed serum substitute medium could pave the way in replacing FBS for BTE studies eliminating the use of controversial FBS and providing a better-defined chemical environment for BTE studies.

**Graphical abstract:** 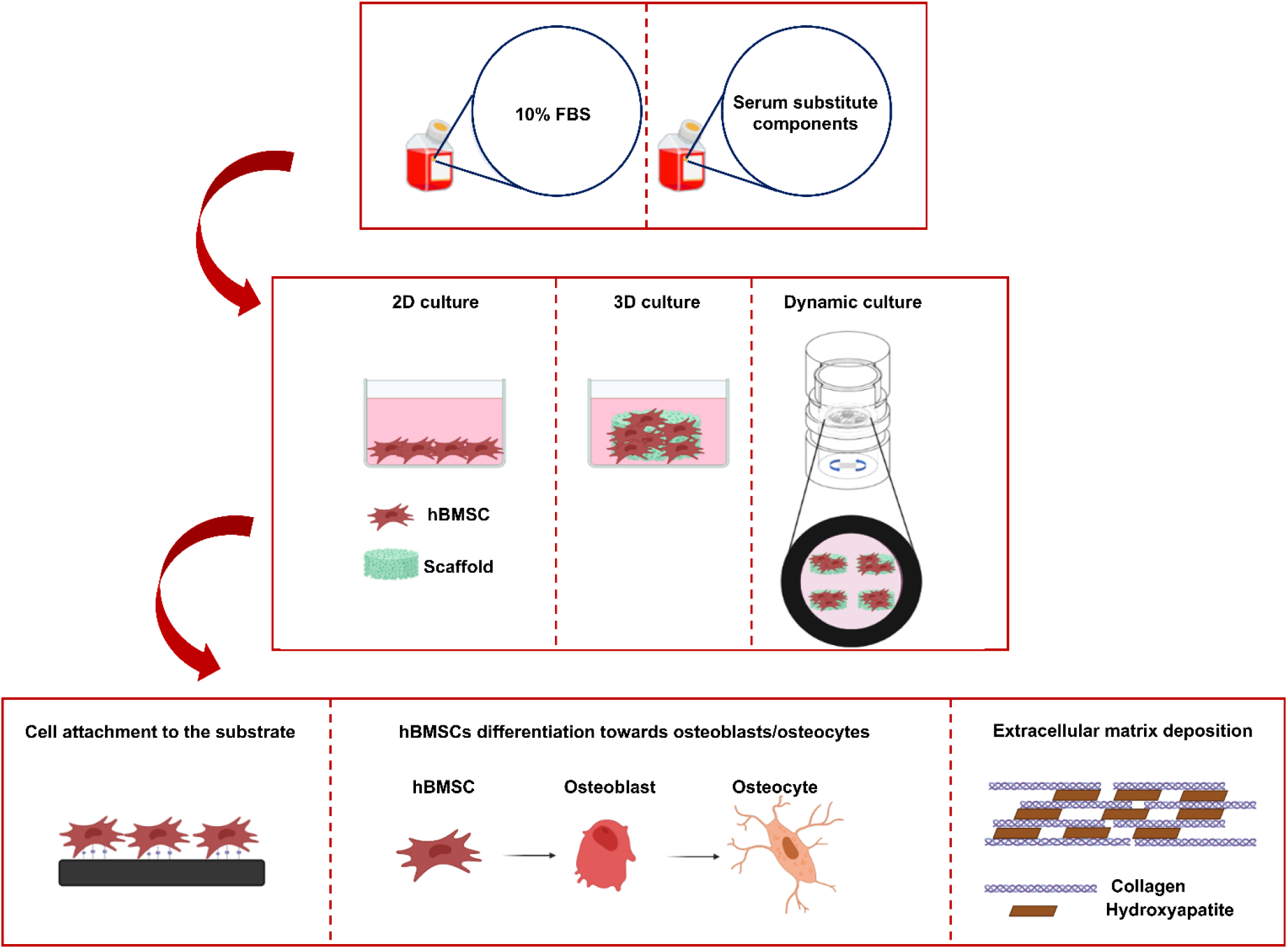

## 1 Introduction

*In vitro* tissue engineering approaches include progenitor cells grown on scaffolds under chemical and/or mechanical stimuli [1]. Many of these *in vitro* methods still make use of fetal bovine serum (FBS). FBS contains hormones, growth factors, attachment factors, protease inhibitors, vitamins, proteins, etc., essential for cell growth and maintenance [2,3]. The exact components of FBS are not known. There have been some reports on certain components within FBS and their concentrations, however, these components/concentrations might differ per FBS batches/brands (Table 1) [3,4]. The variation in composition between different FBS brands and batches impacts the experimental outcomes [5,6]. For instance, an investigation on the elemental component of FBS showed significant elemental variations in different FBS batches which influenced the protein expression of human umbilical vein endothelial cells *in vitro* [6]. Besides having an undefined and complex composition, FBS holds other disadvantages such as safety and ethical issues and shortage in global supply which makes the use of FBS in culture controversial [2]. To overcome the disadvantages of using FBS in culture, chemically defined serum substitute media with known components need to be developed and replace FBS [7].

**Table 1.** Known components of FBS [3]

| Components | Examples of the components |  |
| --- | --- | --- |
| **Proteins** | Serum protein | Albumin, Globulins, α1-Antitrypsin, α2-Macroglobulin |
| | Transport proteins | Transferrin, Transcortin, α1-Lipoprotein, β1-Lipoprotein |
|  | Attachment and spreading factors | Fibronectin, Laminin, serum spreading factor |
| | Enzymes | Lactate dehydrogenase, Alkaline phosphatase, γ-Glutamyl Tranferase, Alanine Aminotransferase, Aspartate Aminotransferase |
| **Hormones** | Insulin, Glucagon, Corticosteroids, Vasopressin, Thyroid Hormones, Parathyroid Hormone, Growth hormone, Pituitary glandotropic factors, Prostaglandins |  |
| **Growth factors** | Epidermal growth factor (EGF), Fibroblast growth factor (FGF), Nerve growth factor (NGF), Endothelial cell growth factor (ECGF), Platelet-derived growth factor (PDGF), Insulin-like growth factors (IGFs), Interleukins, Interferons, Transforming growth factors (TGFs) |  |
| **Fatty acids and lipids** | Free and protein-bound fatty acids, Triglycerides, Phospholipids, Cholesterol, Ethanolamine, Phosphatidylethanolamine |  |
| **Vitamins** | Retinol/Retnoic acid (Vitamin A), Vitamin B-Group, Ascorbic acid (Vitamin C), α-Tocopherol (Vitamin E) | |
| **Trace elements** | Selenium, Iron, Zinc, Copper, Cobalt, Chromium, Iron, etc. |  |
| **Carbohydrates** | Glucose, Galactose, Fructose, Mannose, Ribose, Glycolytic Metabolites |  |
| **Nonprotein Nitrogens** | Urea, Purines/Pyrimidines, Polyamines, Creatinine, Amino acids |  |

Development of serum substitute medium offers several advantages including 1) avoiding the suffering of animals, 2) reducing variability in culture medium composition, 3) supporting cell survival, growth, and homeostasis in the physiological state of a specific tissue or 4) the possibility to recreate pathophysiological states of a specific tissue to investigate potential treatments. The formulation of serum substitute medium depends on many factors such as the cells (primary cells or cell lines), cell culture conditions (mono-culture or co-culture), cell sources (bone marrow, adipose tissue, etc.), applications (cell-based therapy, food industry, or cell-seeded implants), and species (human, canine, etc.) [8–18]. These variations in medium formulations indicate that one universal serum substitute medium might not exist but it should be developed/optimized for each specific cell type and specific application. [7]. Many of these serum substitute medium formulations are already commercially available or can be found in few databases [3,19]. A drawback of commercially available serum substitute media is that their composition is not provided which limits the possibility of studying the influence of soluble factors or drugs on cellular behavior and function.

Bone is a complex multifunctional organ that continuously undergoes a physiological process called bone remodeling to maintain bone strength and mineral homeostasis [20]. This process occurs via balanced activities of its specialized cells, namely, osteoclasts, osteoblasts, and osteocytes [21]. Disturbing the bone remodeling process leads to the development of metabolic bone diseases (e.g. osteoporosis and osteopetrosis) [22]. Bone tissue engineering (BTE) exploits the same tissue engineering principles to develop healthy or pathological human *in vitro* bone models to study bone physiology/pathology or assess drug effects in a pre-clinical setting. FBS has been routinely used to create such *in vitro* models, but to overcome the issues raised by using FBS, a serum substitute medium with known components needs to be formulated for BTE application. The development of a serum substitute medium with known components for BTE has not yet been reported and is the focus of the current study.

In order to use the formulated medium in BTE studies, the newly developed serum substitute medium needs to support the attachment of cells to the substrate, the differentiation of hBMSCs towards osteoblasts/osteocytes, and the deposition of extracellular matrix (ECM). We started developing the serum substitute medium in a step-by-step process by adding the essential components to the medium while the cells were cultured in 2D well-plates and later on 3D silk fibroin scaffolds and evaluated their contribution to cell survival, attachment to the substrate, osteoblast/osteocyte differentiation, collagen production and mineral deposition after 3 weeks of culture (Table 2) [23]. Next, the potential of the newly developed serum substitute medium in supporting cells under application of mechanical loading was assessed. In brief, hBMSCs were seeded on 3D silk fibroin scaffolds in either FBS containing medium or serum substitute medium and placed inside the spinner flask bioreactors. The spinner flask bioreactor generates wall shear stress through a continuous flow of cell culture medium [24]. Cell viability, their attachment to the scaffold, hBMSCs differentiation into osteoblasts/osteocytes, and deposition of mineralized collagenous matrix were investigated after 3 weeks of culture. Such defined serum substitute medium provides the opportunity to study the bone formation process in a controlled environment without the influence of variable components of FBS.

**Table 2.** Serum substitute medium components

| **Component** | **Impact of the components on cells** | **Reference** |
| --- | --- | --- |
| Vitronectin | Enhances attachment of cells to the substrate | [25–27] |
| Human Serum Albumin (HSA) | Acts as antioxidant to prevent oxidative damages to cells through binding to metal ions including copper or iron | [28] |
|  | Binds to various compounds including fatty acids, amino acids, hormones, vitamins, and metal ions (such as Zinc), and transport them to the cells |  |
| antibiotic/antimycotic | Controls the growth of bacterial and fungal contamination | [29] |
| Insulin/transferrin/Selenium | Insulin: enhances glucose and amino acid uptake by cells, intercellular transport, and synthesis of proteins | [15,30] |
|  | Transferrin: transports iron to cells which is a co-factor for enzymes involved in DNA synthesis and prevent extracellular oxidation |  |
|  | Selenium: protects cells from oxidative damages by reducing the production of free radical |  |
| Glutamine | Stable form of glutamine which is an essential amino acid for the synthesis of proteins and nucleic acids | [7,31] |
|  | Acts as a secondary energy source for metabolism |  |
| Non-essential amino acids | Increases collagen synthesis and stabilizes collagen structure | [32–35] |
| Chemically defined lipids | Serves as structural constituents of cell membrane, energy stores, and signaling molecules | [7,36] |
| Basic fibroblast growth factor | Enhances proliferation of mesenchymal stromal cells and stimulate osteoblast differentiation during early stages | [37–39] |
| Bone morphogenetic protein-2 | Stimulates osteogenic differentiation of mesenchymal stromal cells | [37,40,41] |

## 2 Material and methods

### 2-1 Materials

Dulbecco’s modified eagle medium (DMEM high glucose, 41966 and low glucose, 22320), antibiotic/antimycotic (Anti-Anti, 15240062), non-essential amino acids (NEAA, 11140050), and trypsin-EDTA (0.5%, 2530054), Insulin-Transferrin-Selenium (ITS-G, 41400045), GlutaMAX (35050061), chemically defined lipid concentrate (CDlipid, 11905031) were from Life Technologies (The Netherlands). For cell expansion, FBS (F7524) from Sigma-Aldrich (The Netherlands) and for osteogenic differentiation, FBS (SFBS, Lot. No. 51113) from Bovogen (Australia) were used. Basic fibroblast growth factor (b-FGF, 100-18B) was purchased from Peprotech (UK). Recombinant human bone morphogenetic protein-2 (rhBMP-2, 7510200) was purchased from Medtronic Sofamor Danek (USA). Human serum albumin (HSA, A1653) was purchased from Sigma-Aldrich (The Netherlands). Unless noted otherwise, all other substances were of analytical or pharmaceutical grade and obtained from Sigma-Aldrich (The Netherlands).

### 2-2 Medium composition

Cells require nutrients such as amino acids, lipids, carbohydrates, vitamins, trace minerals and inorganic salts to grow, proliferate, and differentiate. Basal media such as Dulbecco’s Modified Eagle’s Medium (DMEM), Roswell Park Memorial Institute (RPMI), and Eagle’s Minimum Essential Medium (EMEM) provide cells with many of these nutrients [29]. To develop a serum substitute medium, the first step is basal medium selection. In this study, we have selected DMEM (low glucose Cat. No. 22320) as the basal medium. This basal medium supplemented with 10% FBS has been shown to be suitable in stimulating osteoblast differentiation of human bone marrow mesenchymal stromal cells (hBMSCs) [23,24,42–44]. DMEM contains a wide variety of amino acids, vitamins, inorganic salts, glucose, and sodium pyruvate (Table 3).

**Table 3.** The components of DMEM and their function on cells

| Components | Impact of the components on cells | references |
| --- | --- | --- |
| Amino acids | Basic building blocks of proteins and essential for cell proliferation and viability | [45] |
| Vitamins | Essential for cell growth and proliferation, act as co-factor for enzymes | [7,46] |
| Inorganic salts | Maintain the osmotic balance and regulate membrane potential which is necessary to transport ions through cell membrane | [47,48] |
| Glucose | Main source of energy for cell metabolism | [29] |
| Sodium Pyruvate | By-product of glucose consumption in the glycolytic pathway and functions as a source of energy | [49] |
| HEPES | Increase buffering capacity of medium to maintain the pH of cultures | [50] |
| Phenol Red | pH indicator | [29] |

#### 2-2-1 FBS containing medium formulation

FBS-containing medium consisted of, 10% FBS (Bovogen), 1% Anti-Anti. To induce osteogenic differentiation of the cells, this medium was supplemented with 50 μg/mL ascorbic-acid-2-phosphate (Sigma-Aldrich, A8960), 100 nM dexamethasone (Sigma-Aldrich, D4902), 10 mM β-glycerophosphate (Sigma-Aldrich, G9422).

#### 2-2-2 Serum substitute medium formulation

The serum substitute medium formulation consisted of DMEM (low glucose, 22320) supplemented with components of table 4. To induce osteogenic differentiation of the cells, this medium was supplemented with 50 μg/mL ascorbic-acid-2-phosphate, 100 nM dexamethasone, 10 mM β-glycerophosphate.

**Table 2.** Concentrations of serum substitute medium components

| Component | Impact of the components on cells | Concentration |
| --- | --- | --- |
| HSA | Acts as antioxidant to prevent oxidative damages to cells through binding to metal ions including copper or iron | 1% w/v |
|  | Binds to various compounds including fatty acids, amino acids, hormones, vitamins, and metal ions (such as Zinc), and transport them to the cells |  |
| Anti-Anti | Controls the growth of bacterial and fungal contamination | 1% v/v |
| ITS-G | Insulin: enhances glucose and amino acid uptake by cells, intercellular transport, and synthesis of proteins | 1% v/v |
|  | Transferrin: transports iron to cells which is a co-factor for enzymes involved in DNA synthesis and prevent extracellular oxidation |  |
|  | Selenium: protects cells from oxidative damages by reducing the production of free radical |  |
| GlutaMax | Stable form of glutamine which is an essential amino acid for the synthesis of proteins and nucleic acids | 1% v/v |
|  | Acts as a secondary energy source for metabolism |  |
| NEAA | Increases collagen synthesis and stabilizes collagen structure | 1% v/v |
| CDlipid | Serves as structural constituents of cell membrane, energy stores, and signaling molecules | 0.1% v/v |
| b-FGF | Enhances proliferation of mesenchymal stromal cells and stimulate osteoblast differentiation during early stages | 10 ng/mL |
| rhBMP-2 | Stimulates osteogenic differentiation of mesenchymal stromal cells | 100 ng/mL |

**Table 3.**
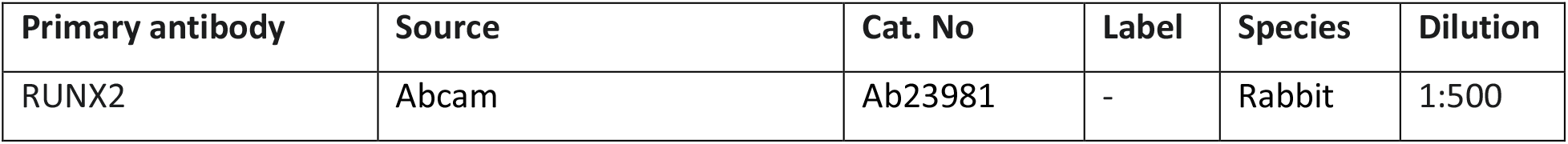
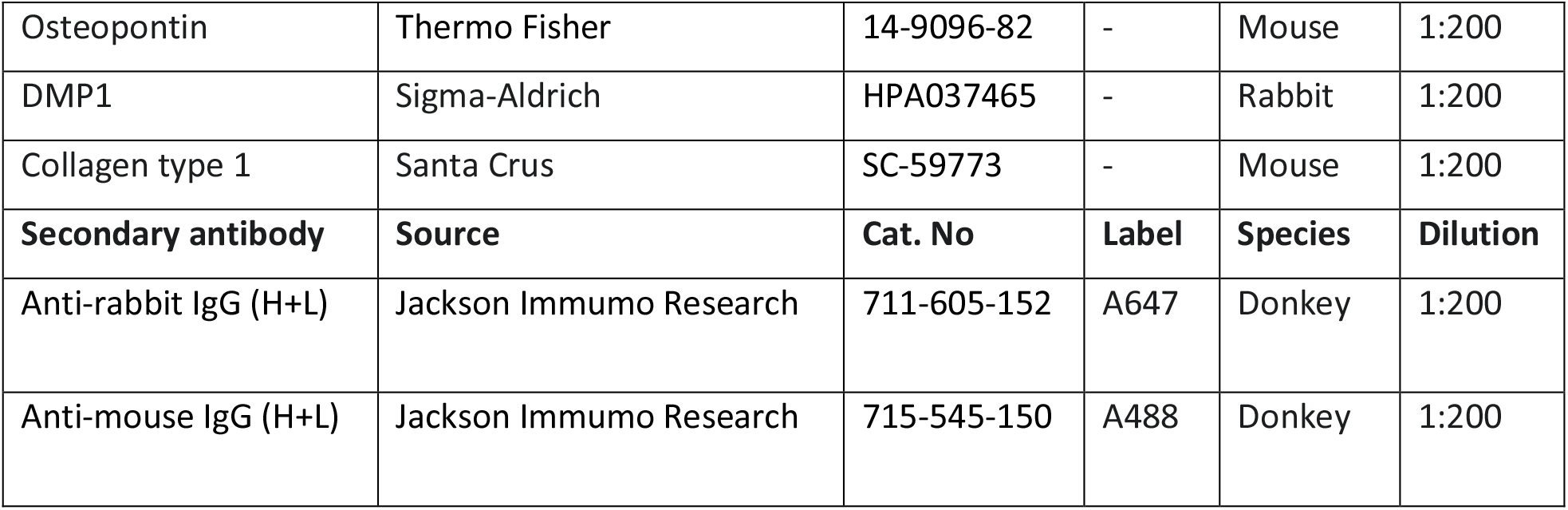
Overview of antibodies used for immunohistochemistry

| Primary antibody | Source | Cat. No | Label | Species | Dilution |
| --- | --- | --- | --- | --- | --- |
| RUNX2 | Abcam | Ab23981 | - | Rabbit | 1:500 |
| Osteopontin | Thermo Fisher | 14-9096-82 | - | Mouse | 1:200 |
| DMP1 | Sigma-Aldrich | HPA037465 | - | Rabbit | 1:200 |
| Collagen type 1 | Santa Crus | SC-59773 | - | Mouse | 1:200 |
| **Secondary antibody** | **Source** | **Cat. No** | **Label** | **Species** | **Dilution** |
| Anti-rabbit IgG (H+L) | Jackson Immuno Research | 711-605-152 | A647 | Donkey | 1:200 |
| Anti-mouse IgG (H+L) | Jackson Immuno Research | 715-545-150 | A488 | Donkey | 1:200 |

### 2-3 Silk fibroin scaffold fabrication

Silk fibroin scaffolds were prepared by cutting and cleaning 3.2 grams of Bombyx mori L. silkworm cocoons (Tajima Shoki., Ltd. Japan) and then degumming them by boiling in 1.5 L UPW containing 0.02 M Na_2_CO_3_ (Sigma-Aldrich, S7795) for 1 hour, whereafter it was rinsed with 10 L cold ultra-pure water (UPW) to extract sericin. Dried purified silk fibroin was dissolved in 9 M lithium bromide (LiBr, Acros organics, 199870025) solution in UPW at 55°C for 1 hour and dialyzed against UPW for 36 hours using SnakeSkin Dialysis tubing (molecular weight cut-off: 3.5 kDa, Thermo Fisher Scientific, The Netherlands). The silk fibroin solution was frozen at −80°C for at least 2 hours and lyophilized (Freezone 2.5, Labconco, USA) for 4 days. 1.7 grams of lyophilized silk fibroin was then dissolved in 10 mL 1,1,1,3,3,3-Hexafluoro-2-propanol (HFIP, Fluorochem, 003409) at room temperature for 5 hours resulting in a 17% (w/v) solution. 1 mL of silk-HFIP solution was added to a Teflon container containing 2.5 grams NaCl with a granule size between 250-300 μm. After 3 hours, HFIP was allowed to evaporate for 4 days. Silk fibroin-NaCl blocks were immersed in 90% (v/v) methanol (Merck, The Netherlands) in UPW for 30 minutes to induce the protein conformational transition to β-sheets and let dry overnight [51]. Scaffolds were cut into disks of 3 mm height with an Accutom-5 (Struer, Type 04946133, Ser. No. 4945193), followed by immersion in UPW for 2 days to extract NaCl. Disc-shaped scaffolds were made with a 5 mm diameter biopsy punch (KAI medical, Japan) and autoclaved in phosphate buffered saline (PBS, Sigma-Aldrich, P4417) at 121°C for 20 minutes.

### 2-4 Cell isolation, expansion, and subsequent seeding

hBMSCs were isolated from 2 donors of human bone marrow (Lonza, USA) and characterized as previously described [52]. Passage 4 hBMSCs were expanded (2500 cell/cm^2^) in expansion medium (DMEM high glucose containing 10% FBS Sigma, 1% Anti-Anti, 1% NEAA, and 1 ng/ml b-FGF) for 9 days, the medium was replaced 3 times per week. At day 9, cells were 80% confluent and trypsinized and proceed as follow.

#### 2-4-1 Cells seeded in well-plates (2D set-up)

The wells of a 48-well-plate (Greiner bio-one, CELLSTAR, 677-180) were coated with 5 μg/ml vitronectin (Peprotech, 140-09) diluted in PBS. Briefly, 100 μL of vitronectin solution was added to each well and incubated in an incubator (37°C, 5% CO_2_) for 2 hours and then at 4°C overnight. The following day, the vitronectin solution was aspirated, the wells were rinsed 3 times with PBS, and the well-plate was pre-warmed in an incubator before seeding cells in wells. 2500 cells were seeded per well resulting in 2500 cells/cm^2^, cultured in either **FBS containing medium** or **Serum substitute medium**, both supplemented with osteogenic differentiation factors (50 μg/mL ascorbic-acid-2-phosphate, 100 nM dexamethasone, 10 mM β-glycerophosphate). The medium was refreshed 3 days a week for 3 weeks.

##### 2-4-1-1 Cell attachment to the well-plate

To determine the amount of DNA as a measure of the number of cells per well after 3 weeks of culture, DNA content was measured. Cell-seeded wells (n=3 per group) were rinsed with PBS and incubated in 500 μL of cell lysis solution containing 0.2% (v/v) Triton-x-100 (Merck, 1.08603.1000) in 5 mM MgCl_2_ (Sigma-Aldrich, M2393) for 30 minutes. Next, the content of the well was transferred into an Eppendorf tube and incubated with 500 μL digestion buffer (500 mM phosphate buffer, 5 mM L-cystein (Sigma-Aldrich, C1276), 5 mM EDTA (Sigma-Aldrich, 1.08421.1000)) containing 140 μg/ml papain (Sigma-Aldrich, P-5306) at 60°C overnight in a water bath shaker (300 RPM). Next, samples were centrifuged at 3000g for 10 minutes and the DNA concentration was measured by Qubit dsDNA HS assay kit (Life Technologies, Q32851). Briefly, 10 μL of samples and standards were mixed thoroughly with DNA buffer containing DNA reagent (1:200) and the DNA concentration was measured with Qubit 2.0 Fluorometer.

##### 2-4-1-2 Immunohistochemistry

After 3 weeks of culture, cell-seeded wells were rinsed with PBS and fixed with 10% neutral buffered formalin for 30 minutes at 4°C. Then, the wells were rinsed 3 times with PBS and covered with 100 μL 0.5% (v/v) Triton-X 100 (Merck, 1.08603.1000) in PBS for 5 minutes to permeabilize cells. Then, the cells were rinsed with PBS and incubated with 5% (v/v) normal donkey serum and 1% (w/v) BSA (Roche, 10.7350.86001) in PBS for 1 hour at room temperature to block the non-specific antibody binding. Wells were then incubated overnight at 4°C with the primary antibody (Table 4). The wells were washed with PBS 3 times and incubated for 1 hour with secondary antibody solution (Table 4). This was followed by 3 times rinsing of the wells with PBS, nuclei were stained with 4′,6-diamidino-2-phenylindole (DAPI (Sigma-Aldrich, D9542), diluted to 0.1 μg/ml in PBS) for 15 minutes. Wells were rinsed with PBS 3 times and then covered with PBS. The expression of proteins was visualized with a Leica TCS SP5X microscope and images were processed with ImageJ (version 1.53f51). Figures were chosen to be representative images per group for all the samples assessed.

##### 2-4-1-3 Alizarin Red staining

After 3 weeks of culture, cell-seeded wells were rinsed with PBS and fixed with 10% neutral buffered formalin for 30 minutes at 4°C. Then, the wells were rinsed 3 times with PBS and covered with 500 μL 2% Alizarin Red solution (Sigma-Aldrich, 05600) diluted in UPW for 15 minutes. Next, the wells were rinsed with UPW and air dried until imaged with a Zeiss Axio Observer Z1. Figures show representative images per group of all the sample assessed.

##### 2-4-1-4 Measurement of deposited calcium in the well-plate

After 3 weeks of culture, cell-seeded wells (n=3 per group) were rinsed with PBS and incubated in 500 μL of 5% trichloroacetic acid (TCA, Sigma-Aldrich, T6399) in UPW for 30 minutes. Next, the content of the well was transferred into an Eppendorf tube and incubated for 48 hours at room temperature. Then, the solids were separated by centrifugation (3000g, 10 min). 5 μL of the supernatant were mixed with 95 μL of calcium working solution (Stanbio Calcium (CPC) LiquiColor® Test, Stanbio Laboratories) and incubated at room temperature for at least 1 minute. In this assay, the calcium ion concentration is measured by the chromogenic complex formed between calcium ions and o-cresolphthalein. Absorbance at 550 nm was measured and calcium concentration was calculated by comparison to standards of known calcium chloride concentrations.

#### 2-4-2 Cells seeded on 3D scaffolds (3D set-up)

Scaffolds were pre-wetted with medium containing FBS or serum substitute which was supplemented with 5 μg/ml vitronectin for 1 hour at 37 °C. Next, the media were removed, and scaffolds were seeded with 1*10^6^ cells each in 20 μL FBS containing medium or serum substitute medium supplemented with 5 μg/ml vitronectin and incubated in an incubator (37°C, 5% CO_2_) overnight. The next day, cell-seeded scaffolds (n=4 per group) were individually placed in wells of a 48-well-plate. Each well was filled with 1 mL of either **FBS containing medium** or **Serum substitute medium**, both supplemented with osteogenic differentiation factors (50 μg/mL ascorbic-acid-2-phosphate, 100 nM dexamethasone, 10 mM β-glycerophosphate). The medium was refreshed 3 days a week for 3 weeks.

##### 2-4-2-1 Live/dead assay

To indicate the amount of living and dead cells at the end of the culture, a live/dead assay was performed. This assay is a two-color fluorescence cell viability assay that is based on the simultaneous determination of living and dead cells. To perform the assay at the end of the culture, scaffolds were rinsed with PBS and incubated in 500 μL of 2 μM Calcein AM (Sigma-Aldrich, 17783) and 4 μM propidium iodide (Invitrogen, P3566) solution prepared in PBS at 37°C, in dark. After 30 minutes, the staining solution was aspirated, and the scaffolds rinsed with PBS. The scaffolds were then imaged with a TCS SP5X confocal microscope (Leica).

##### 2-4-2-2 Cell attachment to the scaffolds and cell number

To determine the number of cells attached to the scaffold, DNA content was measured after 24 hours and 3 weeks of the culture. Scaffolds (n=3 per group) were washed with PBS and each disintegrated in 600 μL of digestion buffer (500 mM phosphate buffer, 5 mM L-cystein, 5 mM EDTA) containing 140 μg/ml papain using steel balls and a Mini-BeadBeater™ (Biospec, USA). Samples were incubated at 60°C overnight in a water bath shaker (300 RPM). Next, the DNA concentration of the sample was measured as explained in the section 2-4-1-1.

##### 2-4-2-3 Immunohistochemistry and histology

###### Sample preparation

After 3 weeks of culture, scaffolds were cut in half, washed with PBS and fixed with 10% neutral buffered formalin over night at 4°C. Then, the scaffolds were dehydrated in graded ethanol solutions and embedded in paraffin, cut into 10 μm thick sections and mounted on Superfrost Plus microscope slides (Thermo Fisher Scientific, The Netherlands). The sections were dewaxed with xylene (VWR, 1330.20.7) (2 times each time for 5 minutes) and rehydrated to water through ethanol solutions (3 times in 100% ethanol each time for 2 minutes, 1 time in 96% ethanol for 2 minutes, 1 time in 70% ethanol for 2 minutes, and finally 2 minutes in UPW). Then, the sections were ready for immunohistochemical and histological staining and Raman spectroscopy.

###### Immunohistochemistry

Sections were incubated in primary and secondary antibodies as explained in section 2-4-1-2. After staining the nuclei with DAPI for 15 minutes, sections were rinsed with PBS 3 times and then mounted on microscope glass slides with Mowiol (Sigma-Aldrich, 81381). The expression of proteins was visualized with a Leica TCS SP5X microscope and images were processed with ImageJ (version 1.53f51). Figures were chosen to be representative images per group for all the samples assessed.

###### Alizarin Red staining

To identify mineral deposition, sections were stained with 2% Alizarin Red solution diluted in UPW for 15 minutes. Then, the sections were washed with UPW and dehydrated with acetone (Boom, 76050006) and acetone-xylene (1:1) solution each for 30 seconds. Next, the sections were cleared in xylene for 5 minutes and mounted in Entellan (Sigma-Aldrich, 1.07960.0500). The samples were imaged with a Zeiss Axio Observer Z1.

###### Picro-Sirius Red staining

To determine collagen production, sections were stained with 0.1% Picro-Sirius Red (Direct red 80, Sigma-Aldrich, CI35872) for 1 hour. Then, the sections were rinsed in 0.1% acidified water and 0.5% acidified water (Acetic acid, Merck, 1.00056.2500) each for 1 minutes. Next, the sections were dehydrated in 70% ethanol for 1 minute, 95% ethanol for 1 minute, and 3 times in 100% ethanol each for 1 minute, and 2 times in xylene for 5 minutes each. Finally, the sections were mounted in Entellan. The samples were imaged with a Zeiss Axio Observer Z1.

###### Hematoxylin and Eosin (H&E)

To localize the cells within the scaffolds, sections were stained with Mayer’s hematoxylin solution (Sigma-Aldrich, MHS16) for 10 minutes. Then, the sections were rinsed in 0.1% acidified water for 1 minute following 5 minutes washing in slow running tap water. Next, the samples were stained in aqueous eosin Y solution (Sigma, HT110-2-16) for 3 minutes followed by washing in slow running tap water for 1 minute. The sections were dehydrated through 30 seconds in 70% ethanol, 30 seconds in 96% ethanol, 3 times in 100% ethanol each for 30 seconds, and 2 times in xylene for 3 minutes. Finally, the sections were mounted in Entellan. The samples were imaged with a Zeiss Axio Observer Z1.

##### 2-4-2-4 Measurement of deposited calcium on cell-seeded scaffolds

After 3 weeks of culture, scaffolds (n=6 per group) were washed with PBS, and each was disintegrated in 500 μl of 5% trichloroacetic acid (TCA) in UPW using steel balls and a Mini-BeadBeater™ (Biospec, USA). After 48 hours of incubation at room temperature, the solids were separated by centrifugation (3000g, 10 min). The calcium concentration was measured as explained in section 2-4-1-4.

##### 2-4-2-5 Raman spectroscopy

Raman measurements were performed using Alpha 300R confocal Raman microscope (WiTec, Germany) on samples that were prepared as described in section 2-4-2-3. Raman imaging of surface areas of 30*30 μm^2^ with a resolution of 1 μm per spectrum of each sample was conducted using a 532 nm excitation laser with a laser power of 10 mW, incubation time of 2s per spectrum, and 50x (NA 0.55) objective. Focus was acquired within the area of the sample using topography correction with manual learning of a 5×5 surface. One plot was cropped due to wrong focus into an area of 30 by 20 mm. Grating was set at 1200 mm^-1^. All acquired spectra were processed using WiTec Project FIVE 5.1.8.64 software (Witec, Ulm). Before analysis, cosmic rays were removed from data with filter size 2 and dynamic factors 6 and background was subtracted with shape size 500. Then, automated True Component Analysis (TCA) was performed to map the location of mineral, collagen and non-mineralized scaffold. The components identified with TCA were then used to graph the spectrum of minerals and collagen. Mineral and collagen peaks were also found within the samples using a filter on the maximum at peaks 961 +/-20, and 1666 +/- 30.

#### 2-4-3 Cells cultured under dynamic condition (Dynamic culture)

The cells were seeded on 3D scaffolds as explained in section 2-4-2. The day after cell incubation with vitronectin solution, cell-seeded scaffolds (n=4 per group) were transferred to spinner flask bioreactors [24]. Each bioreactor contained a magnetic stir bar and was placed on a magnetic stirrer (RTv5, IKA, Germany) at 300 RPM in an incubator (37 °C, 5% CO_2_) [24]. Each bioreactor was filled with 5 mL either **FBS containing medium** or **Serum substitute medium**, both supplemented with osteogenic differentiation factors (50 μg/mL ascorbic-acid-2-phosphate, 100 nM dexamethasone, 10 mM β-glycerophosphate). The medium was refreshed 3 days a week for 3 weeks. After 3 weeks of culture, DNA content on scaffolds and deposited calcium were measured and immunohistochemistry, histology, and Raman spectroscopy were done on the samples the same as explained in section 2-4-2.

### 2-11 Statistics

GraphPad Prism 9.0.2 (GraphPad Software, La Jolla, CA, USA) was used to perform statistical analysis and to prepare the graphs. Due to small sample size (n<12), normality tests have little power to detect whether the samples come from a Gaussian population; therefore, non-parametric tests have been chosen for statistical analysis. To test for differences in DNA and calcium content in 2D set-up (Figure 1A and D, respectively), DNA content in 3D set-ups after 24 hours and 3 weeks (Figure 2A and F, respectively), calcium content in 3D set-up (Figure 4E), DNA content under dynamic condition (Figure 5B), and calcium content under dynamic condition (Figure 7E), an unpaired Mann–Whitney test was performed. Data of DNA content under dynamic condition after 24 hours (Figure 5A) was tested for differences with the non-parametric Kruskal-Wallis with Dunn’s post hoc tests. Differences between groups were considered statistically significant at a level of p < 0.05.

**Figure 1.**
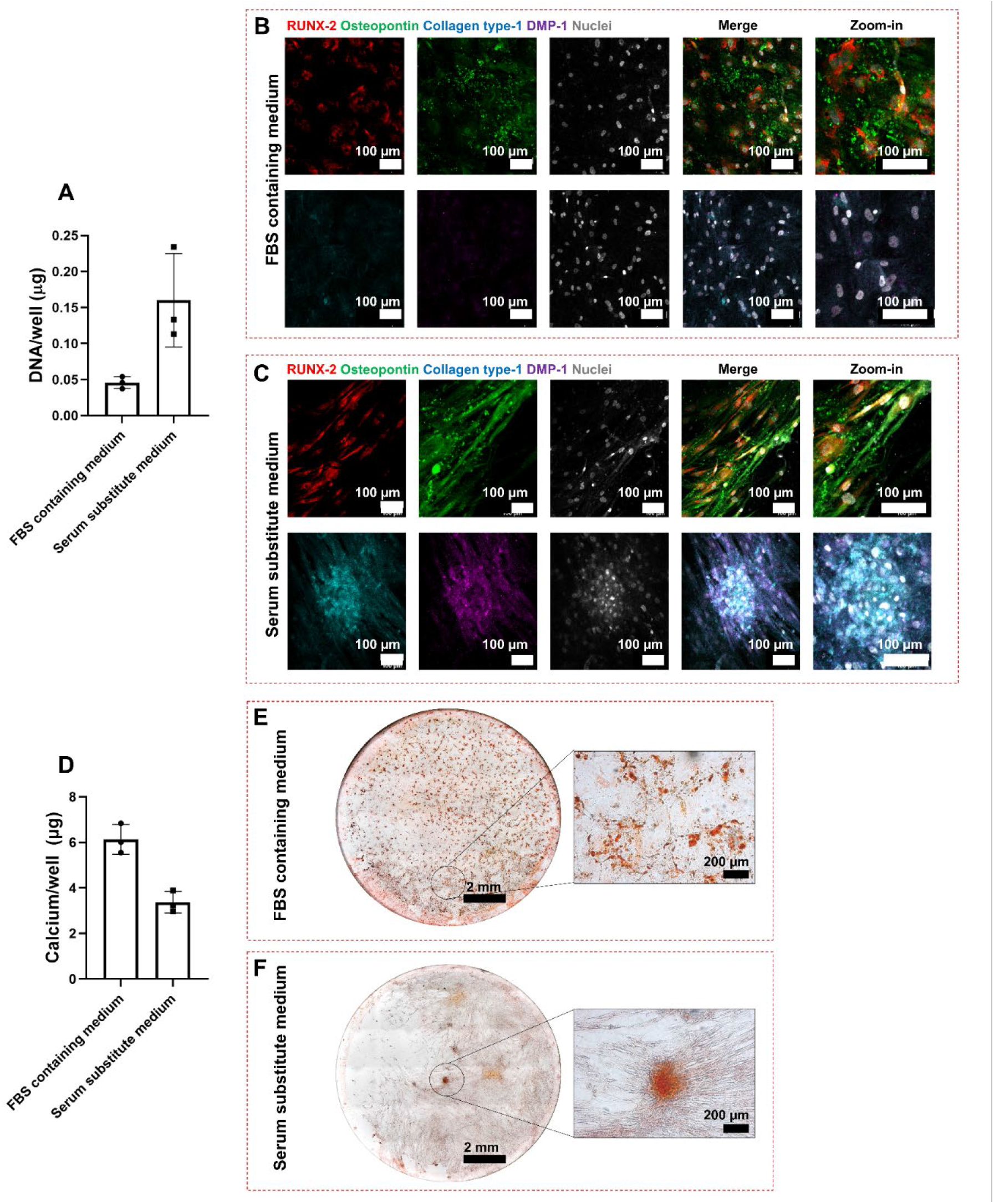
The serum substitute medium supported maintenance of cell attachment to the surface of the well-plate (A). hBMSCs were differentiated towards osteoblasts in FBS containing medium as shown by expression of RUNX-2, collagen type 1, and osteopontin (B). DMP-1 as early osteocyte marker was not expressed by cells cultured in FBS containing medium (B). The serum substitute medium stimulated osteoblast and early osteocyte differentiation of hBMSCs after 3 weeks (C). Calcium was deposited in both FBS containing medium and serum substitute medium. The calcium content in FBS containing medium was higher than in serum substitute medium, even though statistical analysis did not show any significant differences (D). Calcium phosphate deposits were homogenously distributed all over the plate in FBS containing medium (E) while only few mineral nodules were formed in serum substitute medium (F).

**Figure 2.**
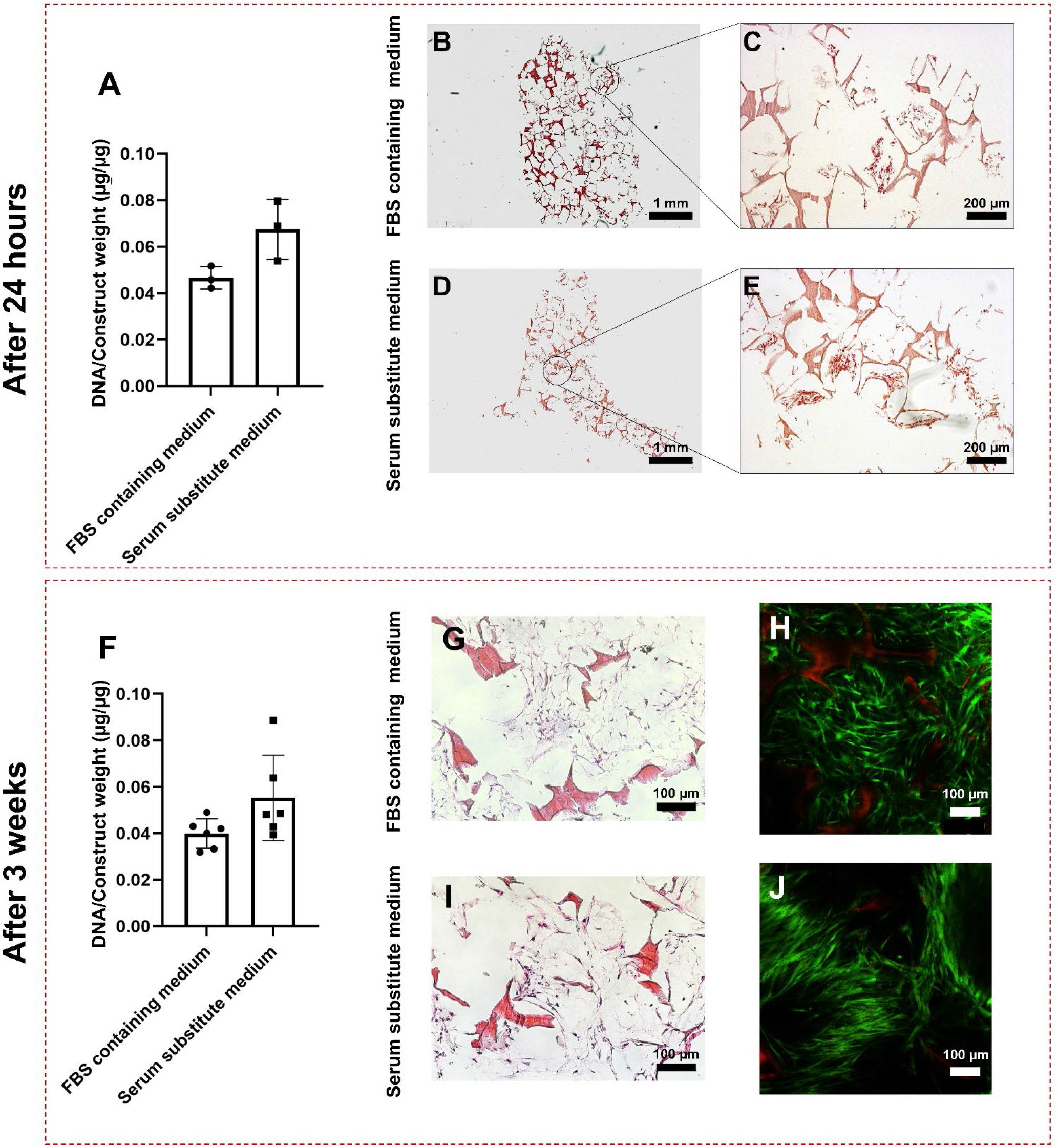
The serum substitute medium supported cells to attach to silk fibroin scaffold: (A) The DNA content of scaffolds after 24 hours of incubation of cells in FBS containing medium or serum substitute medium containing 5 μg/ml vitronectin showed an equal number of cells attached to the scaffolds in both media. The cells were distributed all over the scaffold volume and located between the pores of the scaffold in both FBS containing medium (B-C) and serum substitute medium (D-E). The number of attached cells to the scaffolds was equal in FBS containing medium and serum substitute medium after 3 weeks of culture (F).The cells were located between the pores and attached to the silk fibroin scaffolds in in both media equally (G and I). The cells survived in both media equally after 3 weeks of culture (H and J).

## 3 Results

### 3-1 Serum substitute medium maintained hBMSCs attachment and supported osteoblastic differentiation in a 2D set-up

#### Cell attachment to the substrate

For tissue engineering purposes, anchorage dependent cells should be able to attach to a substrate. FBS has adhesion-promoting properties due to presence of proteins such as fibronectin and vitronectin [26,27]. The serum substitute medium should enable cells to maintain their attachment to the substrate. For the serum substitute medium, vitronectin was chosen as the component to promote cell attachment. The wells of a well-plate were coated with vitronectin, and the cells were seeded in the wells. The maintenance of cell attachment to the surface of well-plates was investigated through measurement of the DNA content of cells attached to the wells. The DNA content of the cells cultured in serum substitute medium showed that the cells maintained their attachment to the surface of the well-plate for 3 weeks (Figure 1A).

#### hBMSCs differentiation towards osteoblasts/osteocytes

After 3 weeks of culturing hBMSCs in either FBS containing medium or serum substitute medium supplemented with osteogenic differentiation factors, the expression of Runt-related transcription factor-2 (RUNX-2) and collagen type 1 as early osteoblast specific markers, osteopontin as a late osteoblast specific protein, and dentin matrix protein-1 (DMP-1) as early osteocyte specific marker was investigated [53,54]. The cells expressed RUNX-2 and osteopontin after 3 weeks of culture in FBS containing medium (Figure 1B) which confirmed the differentiation of hBMSCs towards osteoblasts. As in FBS containing medium, cells expressed RUNX-2 and osteopontin when cultured in serum substitute medium (Figure 1C). RUNX-2 was expressed mostly inside nuclei and cells while osteopontin was found within the ECM. Collagen type 1 was expressed by cells cultured in both FBS containing medium and serum substitute medium (Figure 1B and C). Interestingly, the cells cultured in serum substitute medium formed nodules where collagen type 1 and DMP-1 were highly expressed, while DMP-1 expression was not detected in FBS containing medium cultures.

#### Extracellular matrix deposition

After 3 weeks of culture, the calcium content of wells was measured. Calcium was deposited in both FBS containing medium and serum substitute medium. The calcium content of the cells cultured in FBS containing medium was always higher compared to serum substitute medium, even though statistical analysis did not show any significant differences (Figure 1D). The histological staining of the wells showed that calcium was deposited homogenously over the well in FBS containing medium group, while in serum substitute medium, large mineral nodules were formed on few spots over the well (Figure 1E and F). Thus, formulated serum substitute medium preserved the attachment of hBMSCs to the surface of the well-plate, stimulated hBMSCs differentiation towards osteoblasts/early osteocytes, and supported formation of collagen and mineral nodules.

### 3-2 Serum substitute medium supported hBMSCs attachment and osteoblastic differentiation in a 3D set-up

#### Cell attachment to the substrate

Upscaling to larger 3D BTE constructs requires the use of biomaterial scaffolds that provide a 3D environment to the cells. Serum substitute medium should also be able to promote the attachment of anchorage dependent cells to the 3D scaffolds. In analogy to the 2D study, vitronectin was used as a component to support the attachment of cells to the 3D silk fibroin scaffolds. The attachment of cells to the silk fibroin scaffolds incubated overnight in either FBS containing medium, or serum substitute medium was investigated through measuring the amount of DNA of cells attached to the scaffolds. The number of cells attached to the scaffolds when cultured in serum substitute medium containing 5 μg/mL vitronectin showed no significant differences compared to cultured cells in FBS containing medium (Figure 2A). The Hematoxylin and Eosin (H&E) staining showed the location of seeded cells within the scaffold. In both media, the cells were distributed over the entire scaffold and located within the pores, spanning the void space (Figure 2B-E). These results indicated that the serum substitute medium is able to support cell attachment to the silk fibroin scaffolds. The serum substitute medium supported the maintenance of cell attachment to the silk fibroin scaffolds in serum substitute medium (Figure 2F) which were shown to be present between the pores of the scaffolds (Figure 2G and I). The live/dead assay indicated that the cells were alive in the serum substitute medium after 3 weeks of culture and almost no dead cells were detected (Figure 2H and J).

#### hBMSCs differentiation towards osteoblasts/osteocytes

Cells cultured in the serum substitute medium expressed RUNX-2 and osteopontin similarly as in FBS containing medium (figure 3A and B). As expected, the transcription factor RUNX-2 was detected inside the cells. In FBS containing medium, osteopontin was expressed in the ECM. In contrast, in the serum substitute medium, osteopontin was mostly detected within the cells. The extracellular matrix protein collagen type 1 was expressed in both medium types. DMP-1 as an early osteocyte marker was also expressed within the nuclei and cells in FBS containing medium and serum substitute medium.

**Figure 3.**
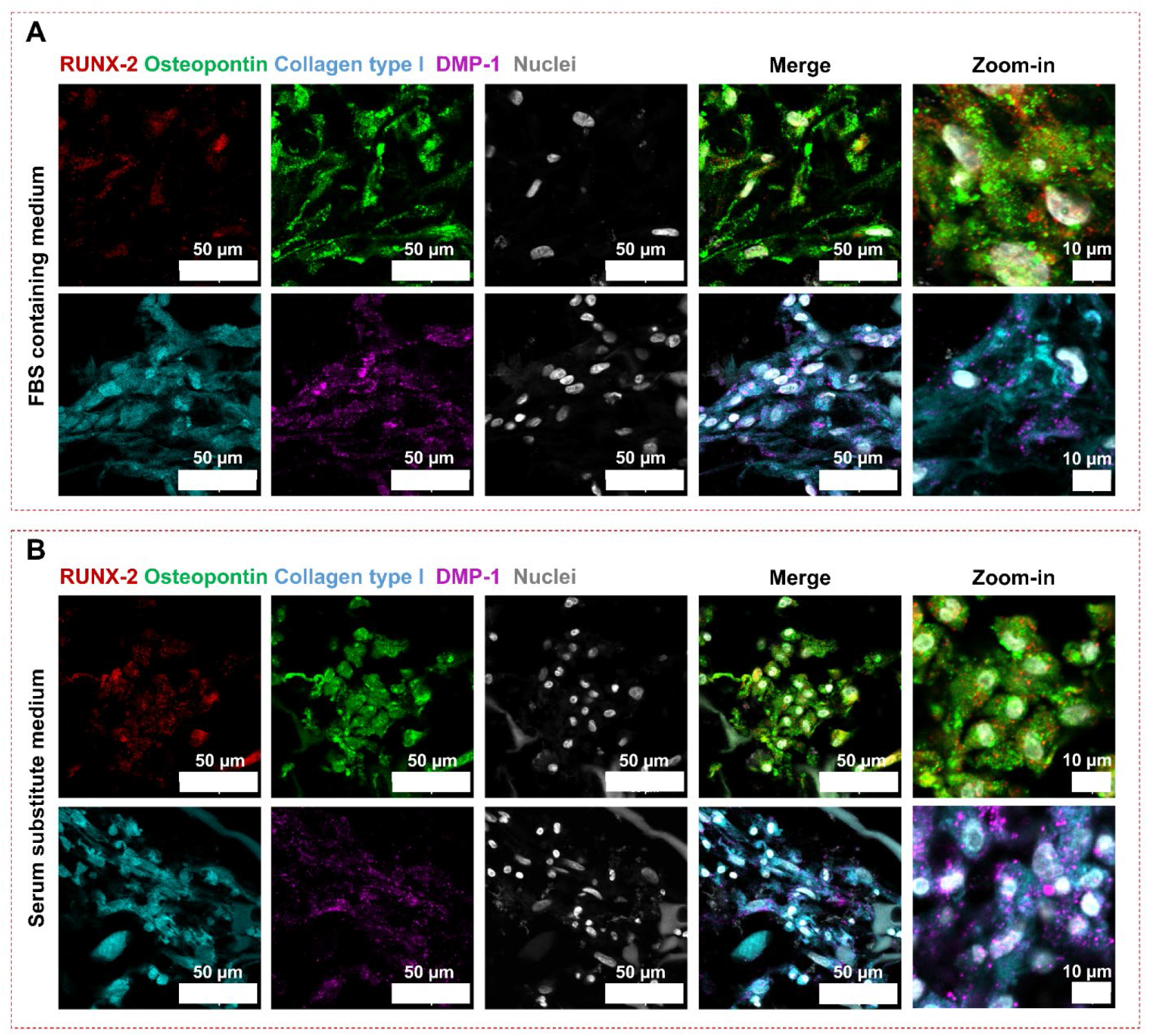
Osteoblast markers expressed by cells cultivated in either medium type. Cells expressed both the early osteoblast differentiation marker RUNX-2 and osteoblast marker osteopontin when cultured in FBS containing medium (A) or in serum substitute medium (B). Collagen, a protein expressed by osteoblasts, was detected in both cultures. DMP-1, an osteocyte-specific protein, was also expressed by cells cultured in FBS containing medium and serum substitute medium.

#### Extracellular matrix deposition

In BTE studies, the medium needs to be able to stimulate cells to express collagen and deposit mineral (mainly carbonated hydroxyapatite), as mineralized collagen forms the basis of bone tissue. Histological staining of samples confirmed the production of collagen by cells cultured in serum substitute medium (Figure 4C) similar as in FBS containing medium (Figure 4A). The produced collagen in serum substitute medium was located in the voids of the pores of the scaffolds as in FBS containing medium. Calcium deposits were detected on cell-seeded scaffolds cultured in FBS containing medium (Figure 4E). Alizarin Red staining showed that these calcium deposits were located on the silk fibroin scaffolds as well as in the ECM produced by cells close to the scaffolds (Figure 4B). However, cells cultured in serum substitute medium did not deposit calcium phosphate on the scaffolds or within the produced ECM (Figure 5E and D). Raman spectroscopy was performed on the cell-seeded scaffolds cultured in FBS containing medium and serum substitute medium. The presence of peaks at approximately 962 cm^-1^, 1299 cm^-1^, and 1441 cm^-1^ representative for phosphate, amide III, and methylated side chains, respectively [55], in FBS containing medium group confirmed the deposited mineralized collagen in this group (Figure 4F). The phosphate and amide peaks were missing in the Raman spectrum of serum substitute medium group (Figure 4F).

**Figure 4.**
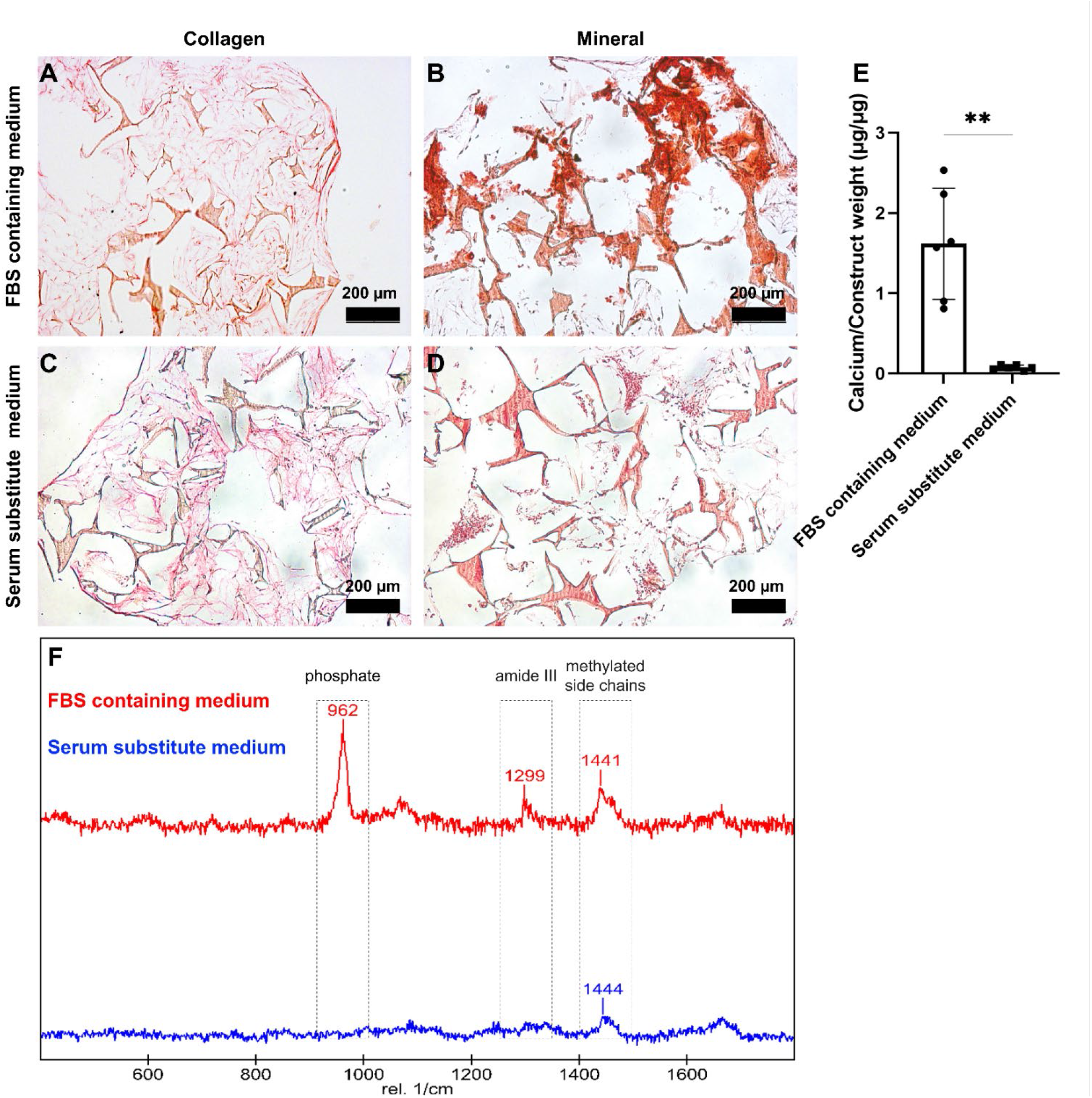
Cells cultured in FBS containing medium produced collagen and showed mineral deposition on scaffolds/ECM (A and B). The cells cultured in the serum substitute medium produced collagen, but no mineral was detected on the scaffolds/ECM (C and D). The calcium assay confirmed the alizarin red staining on the minerals (E). The Raman spectroscopy showed no mineralized collagen formation in serum substitute medium (F).

**Figure 5.**
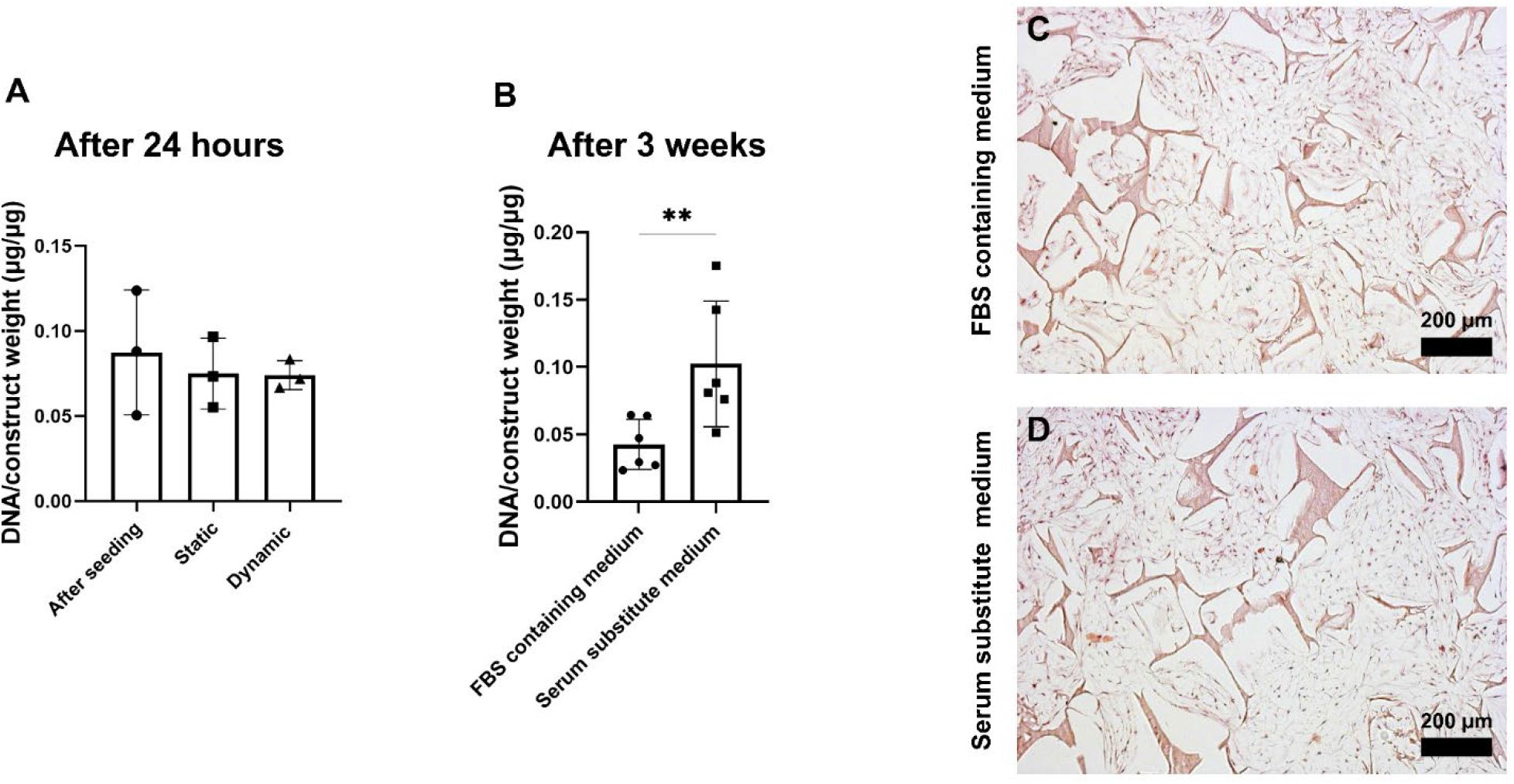
Applied shear stress did not result in cell removal from the scaffolds in serum substitute medium after 24 hours (A). DNA content of scaffolds cultured in FBS containing medium and serum substitute medium for 3 weeks indicated that the cells maintained their attachment to the scaffolds under the dynamic condition (p<0.01) (B). The H&E staining showed that the cells were distributed between the pores of the scaffold in FBS containing medium (C) and in serum substitute medium (D).

### 3-3 Serum substitute medium supported hBMSCs attachment and osteoblastic differentiation in dynamic conditions

#### Cell attachment to the substrate

In order to make sure that cells can form a tissue and investigate the cellular behavior in dynamic conditions, cells need to maintain their attachment to the substrate when they are exposed to mechanical loading. Due to the adhesion-promoting properties of FBS, cells can maintain their attachment to the scaffolds when cultured in FBS containing medium. Serum substitute medium also needs to be able to preserve the attachment of cells to the scaffolds when subjected to mechanical loading such as shear stress. To investigate whether shear stress could result in detachment of cells from the scaffolds in the serum substitute medium, cell-seeded scaffolds were kept in either static or dynamic condition for 24 hours after cell seeding. The number of cells did not change after 24 hours of applied shear stress (Figure 5A). The DNA content of cell-seeded scaffolds was also measured after 3 weeks of culture in either FBS containing medium or serum substitute medium under dynamic condition. The results indicated that the shear stress applied in the dynamic condition did not lift off the cells in serum substitute medium, however the number of cells cultured in serum substitute medium was significantly higher than cells cultured in FBS containing medium (Figure 5B). H&E staining confirmed the presence of the cells in the scaffolds (Figure 5C and D). Moreover, it showed that the cells were well distributed in the scaffolds in both FBS containing medium (Figure 5C) and in serum substitute medium (Figure 5D).

#### hBMSCs differentiation towards osteoblasts/osteocytes

hBMSCs were cultured under continuous shear stress in both media supplemented with osteogenic differentiation factors for 3 weeks. Cells cultured in FBS containing medium expressed RUNX-2, osteopontin, collagen type I, and DMP-1 after 3 weeks of culture which indicated the differentiation of hBMSCs towards osteoblasts and early osteocytes (Figure 6A). hBMSCs also differentiated towards osteoblasts in serum substitute medium as shown by expression of RUNX-2 and osteopontin which were detected inside nuclei and in the ECM, respectively (Figure 6B). Expression of DMP-1 by cells embedded within the collagenous matrix indicated early osteocyte differentiation after 3 weeks of culture in serum substitute medium under mechanical loading (Figure 6B).

**Figure 6.**
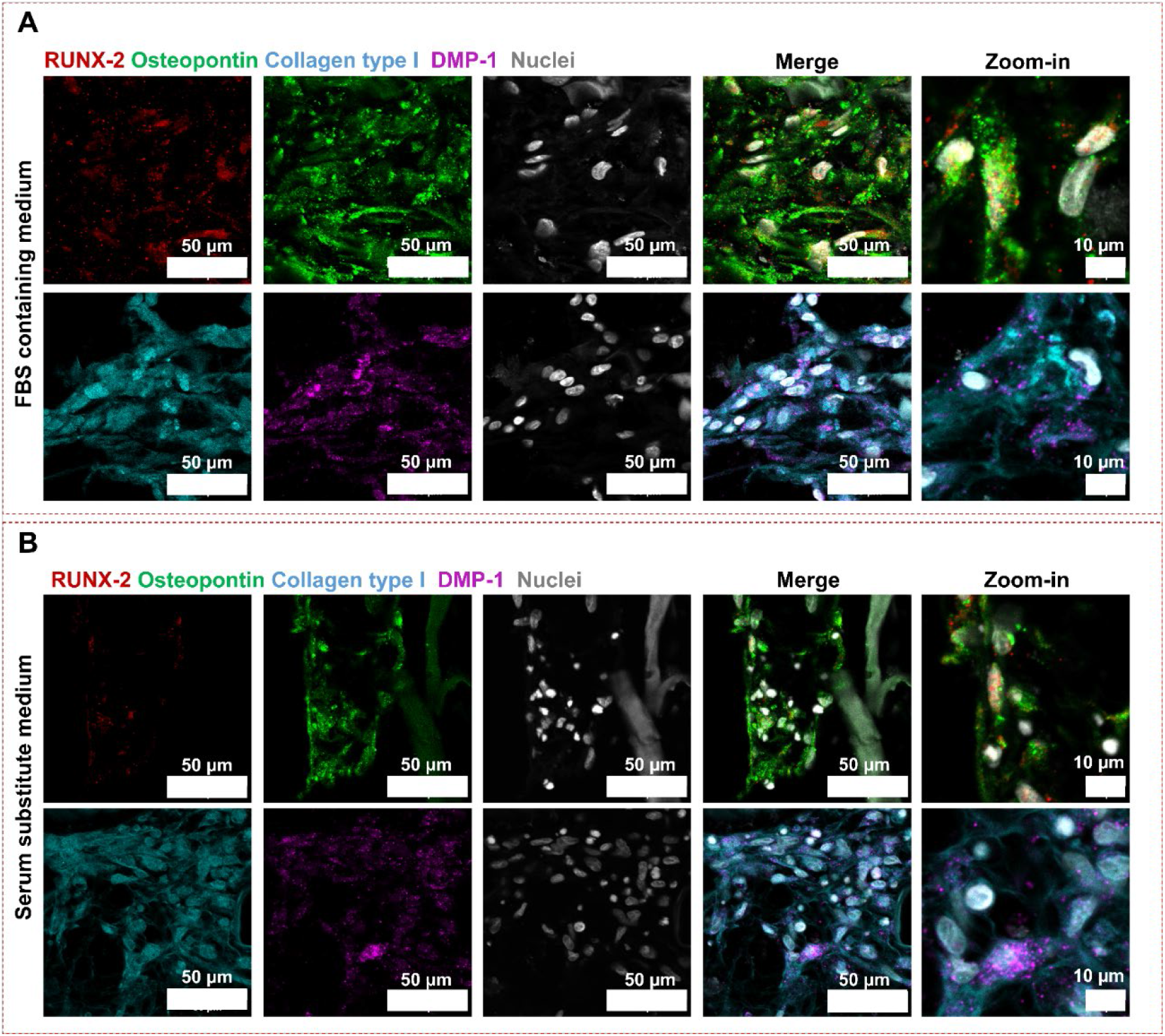
Osteoblast/osteocyte markers expressed by cells cultivated in either medium type under continuous shear stress. Cells expressed both the early differentiation marker RUNX-2 and osteoblast marker osteopontin when cultured in FBS containing medium (A) or in serum substitute medium (B). Collagen type I, a protein expressed by osteoblasts, was detected in abundance in both cultures. DMP-1, an osteocyte-specific protein was detected in both cultures.

#### Extracellular matrix deposition

After 3 weeks of culturing hBMSCs under dynamic conditions in either medium, collagen was expressed as confirmed by histological images (Figure 7A and C). The collagen was laid down between the pores of the scaffolds in both media conditions. The expressed collagen by cells cultured in serum substitute medium seemed denser between the scaffold pores compared to FBS containing medium. Under dynamic conditions, the amount of deposited calcium in serum substitute medium was equal to the FBS containing medium group (Figure 7E). The deposited calcium on the scaffolds and/or ECM was enhanced in the serum substitute medium group under dynamic condition compared to 3D culture in static condition where no mineral was formed (Figure 4E and 7E). Alizarin red staining indicated that in the FBS containing medium group, minerals were deposited on/within the silk fibroin scaffold and ECM, while, in the serum substitute medium, the minerals were found between the pores of the scaffolds (Figure 7B and D) where collagen was also deposited (Figure 7C). Raman spectroscopy was performed on the cell-seeded scaffolds cultured in FBS containing medium and serum substitute medium under dynamic conditions. The presence of the peaks of phosphate (approximately 960 cm^-1^), amide III (1240-1320 cm^-1^), methylene side chains (CH_2_ at 1450 cm^-1^), and amide I (1616-1720 cm^-1^) demonstrated the formation of mineralized collagenous matrix in serum substitute medium (Figure 7F).

**Figure 7.**
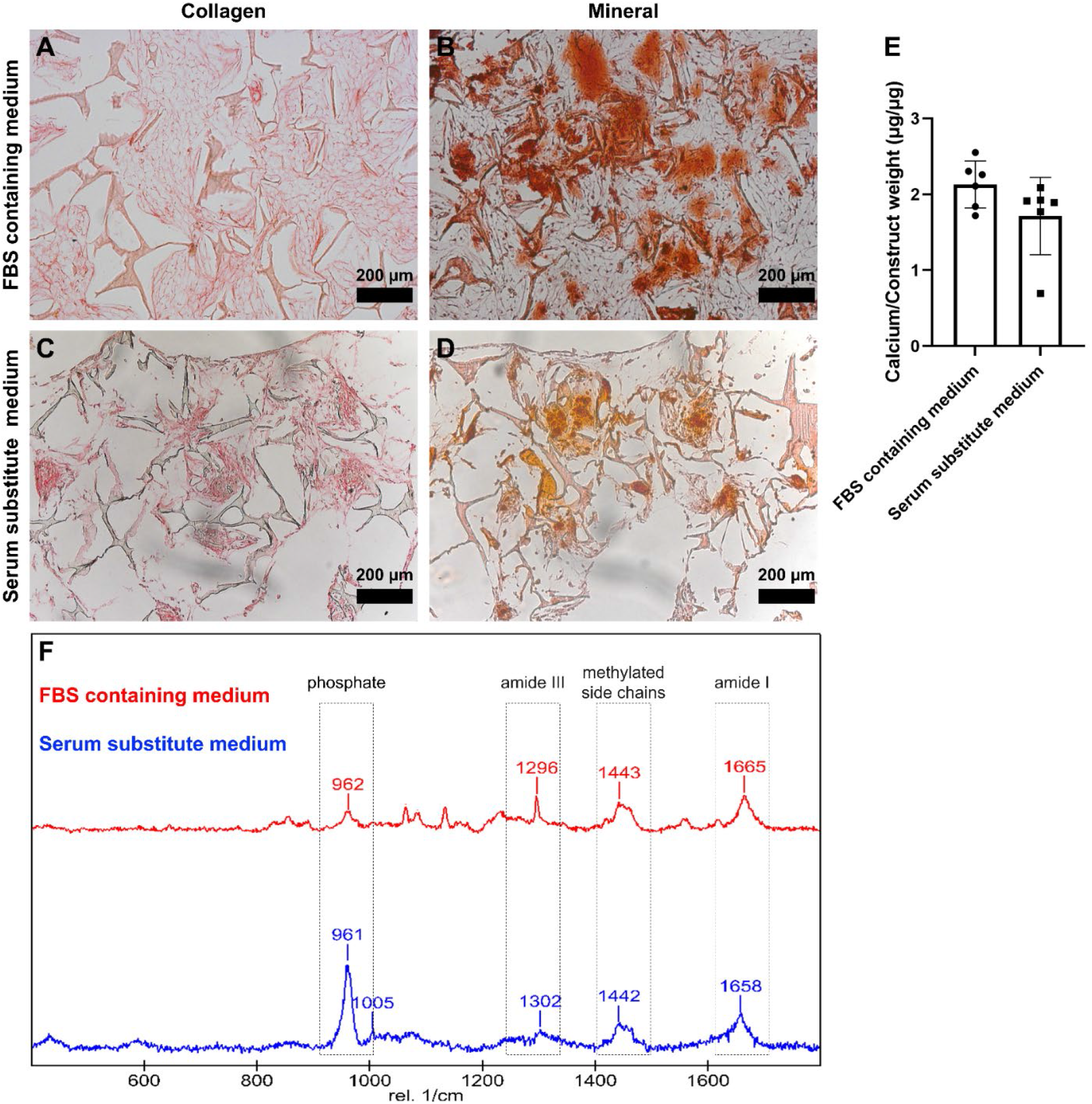
Cells cultured in FBS containing medium and serum substitute medium produced collagen and showed mineral deposition on scaffolds/ECM (A-D) under dynamic condition. Equal amount of calcium was detected in both FBS containing medium and serum substitute medium (E). The Raman spectroscopy showed mineralized collagen formation in both groups (F).

## Discussion

FBS is still routinely used as a cell culture supplements in many research lines despite its known disadvantages. The undefined, complex, and inconsistent composition of FBS has been shown to greatly affect the experimental outcomes and to contribute towards low reproducibility of data [2,56]. Thus, FBS needs to be replaced, ideally by a chemically defined medium. Due to the differences in cells’ requirements in growth, the defined media need to be optimized for each specific cell type and application. In the present study, we developed a serum substitute medium in a step-by-step approach for BTE studies. The potential of this medium in hBMSCs attachment to the substrate, their differentiation towards osteoblasts/osteocytes, and production of a mineralized collagenous matrix in 2D and 3D substrates and under the application of mechanical loading was evaluated.

In the present study, the developed serum substitute medium stimulated hBMSCs to differentiate into osteoblasts as shown by expression of RUNX-2 and osteopontin when the cells where cultured in 2D well-plates and on 3D silk fibroin scaffolds with or without the application of mechanical loading. RUNX-2 is a transcription factor which is expressed at early stages of osteoblast differentiation and regulates the expression of several specific genes related to osteoblasts [57]. The ECM protein osteopontin is expressed by mature osteoblasts and it is believed that it induces mineralization of collagenous fibrils through crystal growth inhabitation in ECM [58,59]. The components of serum substitute medium also supported cells to differentiate further into early osteocytes shown by expression of DMP-1. FBS has been shown to be a cocktail of growth factors including b-FGF and BMPs which greatly influence cell growth and differentiation [56,60]. b-FGF and BMP-2 were chosen as two specific growth factors to stimulate osteoblast differentiation of hBMSCs. b-FGF is a potent mitogen that enhances MSCs proliferation and maintains their differentiation potential [61]. BMP-2 is a member of the transforming growth factor-β superfamily that is well-known for its potential in osteoblast differentiation of MSCs and bone formation *in vitro* and *in vivo* [62]. Without any growth factors, hBMSCs could not differentiate towards osteoblasts (SI 2C) and the addition of b-FGF to the serum substitute medium seemed to be essential to induce osteogenic differentiation of hBMSCs (SI 2D). The presence of BMP-2 alone in serum substitute medium negatively influenced the number of cells (SI 2E), while the addition of BMP-2 to the serum substitute containing b-FGF did not show any significant influence on osteogenic differentiation of hBMSCs (SI 2F). Previous studies on the influence of combination of b-FGF and BMP-2 on osteoblast differentiation of MSCs and bone formation showed that b-FGF had a proliferative role during early stages of differentiation while BMP-2 promoted osteogenic differentiation in the later stages [37,63,64]. It seemed like, in serum substitute medium, b-FGF was needed to induce cell proliferation which further differentiate towards osteoblast in the presence of osteogenic differentiation supplements (dexamethasone, ascorbic acid, and β-glycerophosphate) and the addition of BMP-2 was not needed.

Collagen is the most prominent constituent of the bone organic matrix which is synthesized by osteoblasts. Glycine and proline are among the amino acids that form the collagen structure and their insufficient availability could result in lack of collagen synthesis. [34,65]. Influence of proline on collagen synthesis by fibroblasts as the main collagen synthesized cells has been shown before. The exogenous proline upregulated collagen expression by fibroblasts in the absence of glutamine [34]. Another study indicated the collagen synthesis, mainly collagen type 2, by chondrocytes has improved in the presence of glycine. Basal medium (DMEM) already contains several amino acids such as glycine, however, the addition of 1% v/v NEAA containing 10 mM glycine and 10 mM proline to the serum substitute medium was able to improve ECM production by hBMSCs during osteogenic differentiation (SI 3).

The secreted collagen by osteoblasts gets mineralized during bone formation. To mineralize bone, osteoblasts express membrane-bound alkaline phosphatase (ALP) or secreted matrix vesicles containing alkaline phosphatase (ALP) which cleaves phosphate sources (*i*.*e*., pyrophosphate). The composition of free phosphate with calcium results in calcium phosphate deposition within ECM [66]. FBS has shown to induce mineralization of collagenous matrix which could be attributed to its ALP activity [67,68]. The deposition of minerals in FBS containing medium in 2D cultures and 3D cultures could be influenced by both cellular and FBS ALP activity. In 2D cultures, the homogenous distribution of deposited minerals in FBS containing medium could be attributed to the alkaline phosphatase (ALP) activity of FBS which could provide free phosphate to the culture and thus calcium phosphate deposited all over the well-plate. While in serum substitute medium, hBMSCs needed to differentiate into osteoblasts which express ALP. These osteoblasts cleave β-glycerophosphate resulted in increasing free phosphate to the medium and deposition of calcium phosphate only in locations where cells have differentiated into osteoblasts.

Interestingly, mineral deposition did not take place in cells in 3D set-ups cultured in serum substitute medium unlike in 2D culture. The differences in matrix mineralization in 2D and 3D cultures could be due to less cell-cell contact in 3D cultures compared to 2D cultures. The 3D porous structure of silk fibroin scaffolds might influence the cell-cell interaction which resulted in lack of mineral nodules formation. Previous studies have shown the direct influence of cell density with ECM formation/mineralization during osteoblast differentiation [69,70]. An increase in cell density might enhance the cell-cell interactions and increase the deposition of collagenous matrix. Thus, it would be good to investigate the influence of a higher cell density on matrix mineralization. To increase calcium phosphate deposition, serum substitute medium was supplemented with either vitamin D (10 nM) or high concentration of rhBMP-2 (1000 ng/ml). Vitamin D showed no effect on calcium phosphate deposition despite what was expected [71,72]. The high/non-physiological dose of rhBMP-2 is able to push cells to deposit calcium phosphate, while at the same time it inhibited the collagenous matrix production (SI 4). Moreover, stimulating cells with high doses of growth factors might potentially disguise the influence of other soluble factors on cells as the combination of growth factors could have synergistic effects on cells [37,73,74].

Bone is continuously subjected to different mechanical forces such as shear stress, hydrostatic pressure, and mechanical stretch and tension due to body movement [75]. Early *in vivo* studies have shown that mechanical loading stimulates the formation of woven bone [76–80]. Woven bone forms during skeletal development or fracture healing where a rapid pace of matrix deposition is needed [77]. Woven bone formation is triggered under a variety of loading conditions where pre-osteoblasts lay down randomly oriented collagen that becomes highly mineralized [80,81]. In the current study, shear stress has been applied to hBMSCs seeded on silk fibroin scaffolds. The application of shear stress increased mineral formation in FBS containing medium and induced mineralization in the serum substitute medium compared to static condition. The serum substitute medium without the presence of components such as ALP which could influence mineralization process during *in vitro* bone formation indicated the crucial role of mechanical loading in deposition of woven bone *in vitro*.

The current study obviously has its limitations. For example, in the present study, FBS has been used as the gold standard/control meaning that the outcomes of the developed serum substitute medium were compared to the FBS containing medium. This comparison raises the question of how to validate the results if the outcomes of the control group could be unreliable due to the variable nature of FBS. It should also be noted that in the current study, the focus was on development of a serum substitute medium for osteogenic differentiation of hBMSCs and the isolation and expansion of hBMSCs were still done in FBS containing medium. Exposure of cells to FBS during isolation and expansion steps might affect the cell phenotype and their differentiation capacity. A recent study showed that BMSCs lost their chondrogenic potential after expansion in FBS containing medium [82]. Thus, the serum substitute medium for hBMSCs isolation and expansion also needs to be optimized to have fully FBS-free studies. As every component within the serum substitute medium is known, in future studies, different components can be added to this medium to systematically investigate their influence on cells in *in vitro* bone formation, allowing to extract influential parameters in a more precise way.

## Conclusion

In this study we developed a serum substitute medium able to replace FBS in BTE studies. The developed serum replacement medium supported hBMSCs attachment, differentiation and extracellular matrix production both in 2D and 3D. The most prominent difference was that for mineralization in 3D, mechanical stimulation was essential and representative of mineralization process during woven bone formation. The serum substitute medium has the potential to eliminate the use of FBS in the creation of *in vitro* bone models using BTE approaches and provides an opportunity to systematically study the influence of soluble factors on the bone formation in a less variable environment and without being overshadowed by unknown factors within FBS.

## Acknowledgement

This work has been financially supported by the Dutch Ministry of Education, Culture and Science (Gravitation Program 024.003.013). We would like to thank Dewy van der Valk for her help in Raman spectroscopy measurement and analysis.

