## Supplementary figures for "Development of serum substitute medium for bone tissue engineering"

**SI 1:** Vitronectin was used as a component to promote the attachment of cells to the 3D silk fibroin scaffolds. Cells were incubated in either FBS containing medium or serum substitute medium with or without 5 µg/ml vitronectin. The attachment of cells to the silk fibroin scaffolds incubated overnight in either FBS containing medium or serum substitute medium was investigated through measuring the amount of DNA of cells attached to the scaffolds. The presence of 5 µg/ml vitronectin in serum substitute medium promoted the attachment of cells to the substrate (Figure SI 1A). After 3 weeks of culture, H&E staining showed the distribution of cells between the pores of scaffolds in FBS containing medium (Figure SI 1B) and serum substitute medium containing 5 µg/ml vitronectin (Figure SI 1D). The cells incubated in serum substitute medium without vitronectin distributed sporadically between scaffold pores (Figure SI 1C).

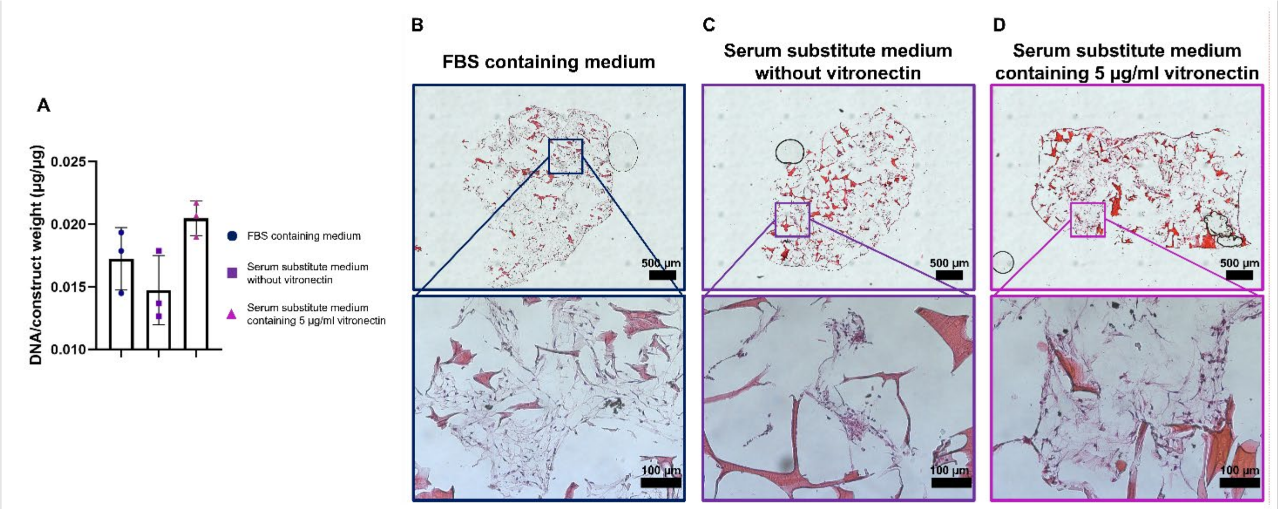


*Figure S1. The DNA content of scaffolds after 24 hours of incubation showed that the serum substitute medium containing 5 µg/ml vitronectin increased attachment of cells to silk fibroin scaffold (A). H&E staining showed that after 3 weeks of culturing cells in FBS containing medium and serum substitute medium containing 5 µg/ml vitronectin distributed all over the scaffolds (B and D). While in serum substitute medium without vitronectin, cells sporadically spread between scaffold pores (C).*

**SI 2:** Without any growth factors, hBMSCs could not differentiate towards osteoblasts (SI 2C) and the addition of b-FGF to the serum substitute medium seemed to be essential to induce osteogenic differentiation of hBMSCs (SI 2D). The presence of BMP-2 alone in serum substitute medium negatively influenced the number of cells (SI 2A and E), while the addition of BMP-2 to the serum substitute containing b-FGF did not show any significant influence on osteogenic differentiation of hBMSCs (SI 2F).

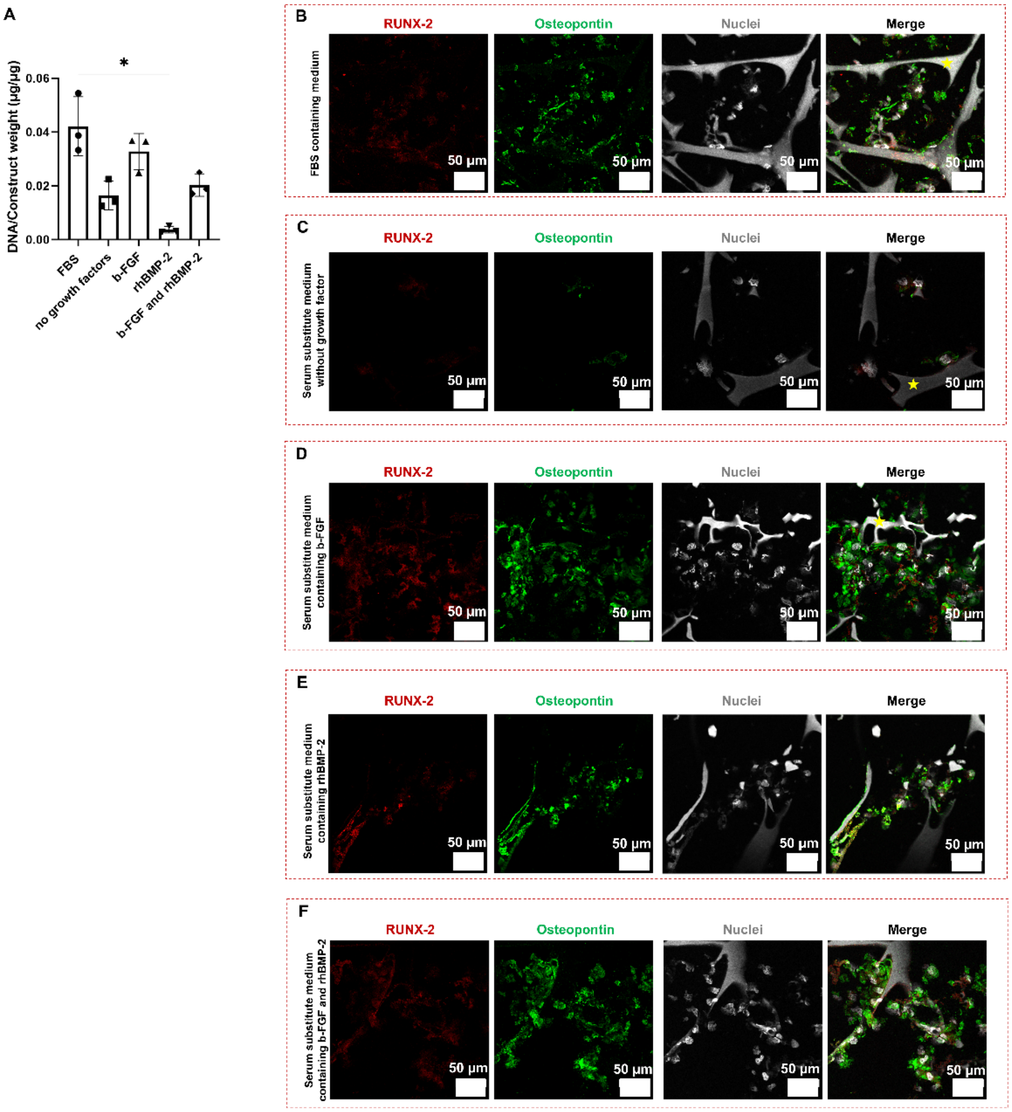


*Figure S2. hBMSCs could differentiate towards osteoblasts as shown by the expression of RUNX-2 and osteopontin in FSB containing medium (B). Lack of growth factors did not induce osteoblast differentiation in hBMSCs (C). The presence of b-FGF was needed for osteogenic differentiation in serum substitute medium (D). The addition of BMP-2 alone to serum substitute medium negatively affected the number of cells (A), while in combination of b-FGF did not show any significant changes in osteogenic differentiation of hBMSC.*

**SI 3:** Collagen is the most prominent constituent of the bone organic matrix which is synthesized by osteoblasts. Glycine and proline are among the amino acids that form the collagen structure and their insufficient availability could result in lack of collagen synthesis. Influence of proline on collagen synthesis by fibroblasts as the main collagen synthesized cells has been shown before. In this study, exogenous proline upregulated collagen expression by fibroblasts in the absence of glutamine. Another study indicated the collagen synthesis, mainly collagen type 2, by chondrocytes has improved in the presence of glycine. Basal medium (DMEM) already contains several amino acids such as glycine, however, the addition of 1% v/v NEAA containing 10 mM glycine and 10 mM proline to the serum substitute medium was able to improve ECM production by hBMSCs during osteogenic differentiation (SI 3).

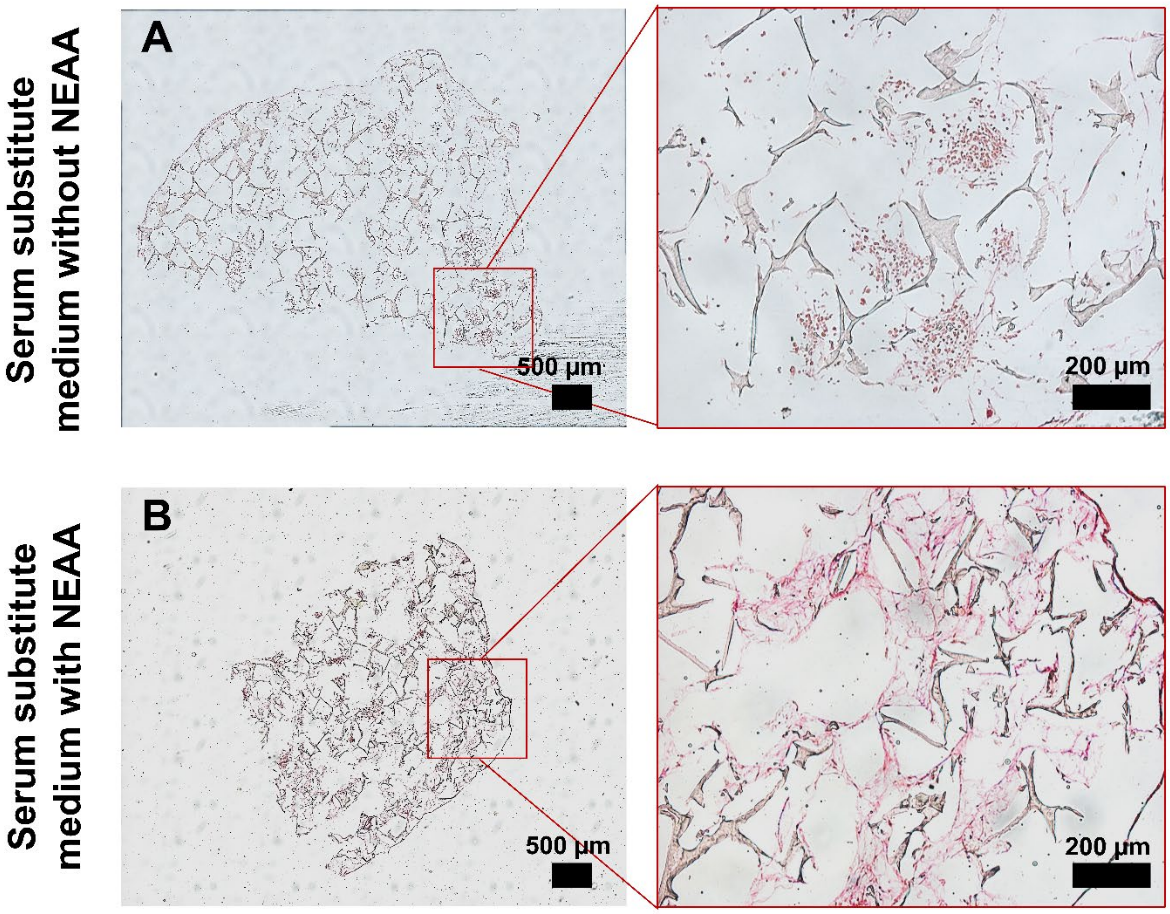


*Figure S3. Exogenous NEAA containing glycine and proline enhanced collagen formation in the serum substitute medium.*

**SI 4:** To increase calcium phosphate deposition, serum substitute medium was supplemented with either vitamin D (10 nM) or high concentration of rhBMP-2 (1000 ng/ml). Vitamin D showed no effect on calcium phosphate deposition despite what was expected. The high/non-physiological dose of rhBMP-2 is able to push cells to deposit calcium phosphate, while at the same time it inhibited the collagenous matrix production (SI 4).

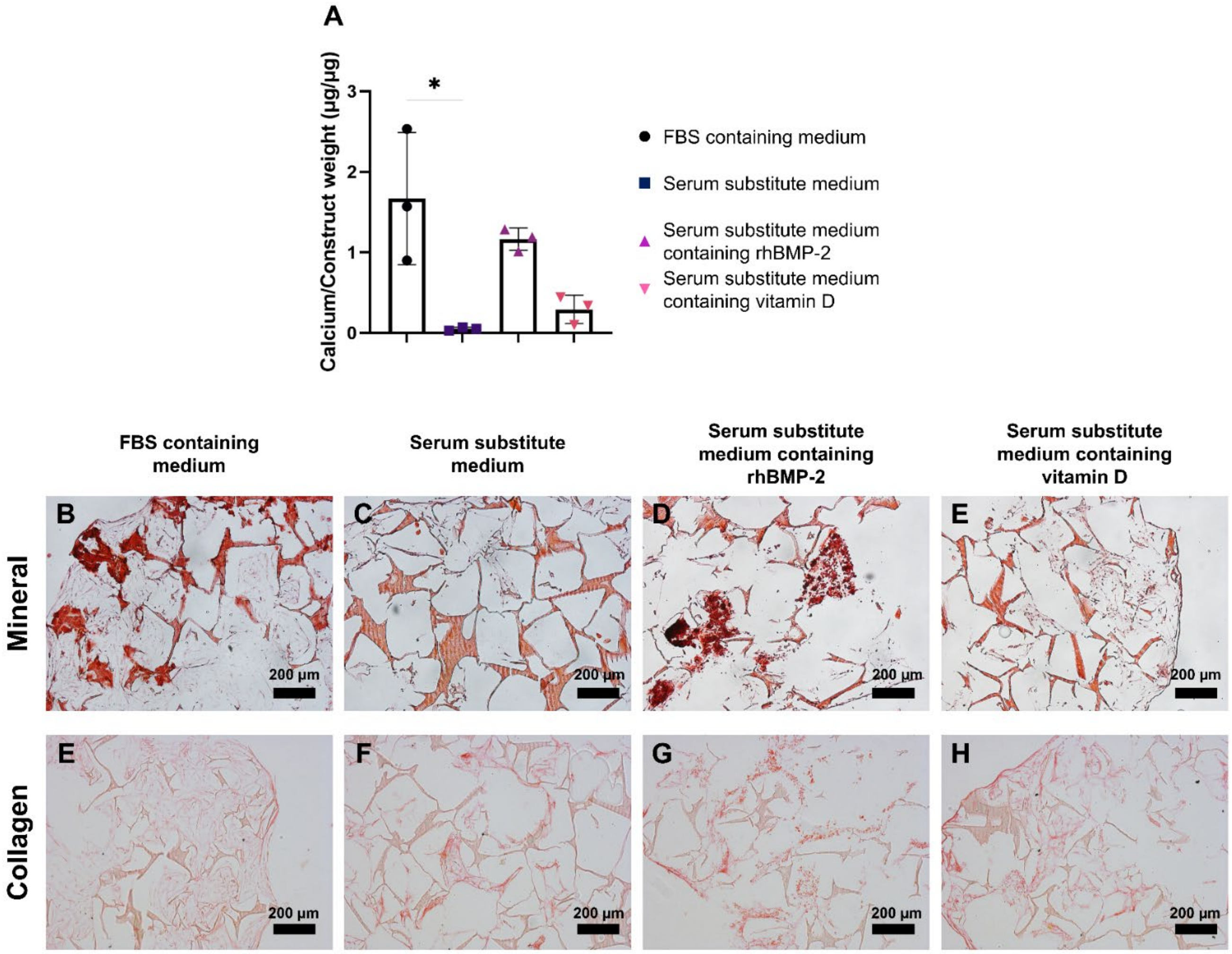


*Figure S4. Addition of vitamin D to serum substitute medium did not have any influence of collagen production and mineral deposition (A, E, H). Addition of high dosage of rhBMP-2 to serum substitute medium increased mineral deposition (A and D) while negatively influenced collagen production (G).*
